# Structure of ATP synthase from ESKAPE pathogen *Acinetobacter baumannii*

**DOI:** 10.1101/2021.07.10.451757

**Authors:** Julius K Demmer, Ben P Phillips, O Lisa Uhrig, Alain Filloux, Luke P Allsopp, Maike Bublitz, Thomas Meier

## Abstract

*Acinetobacter baumannii* is a clinically relevant pathogen which causes multi-drug resistant, hospital-acquired infections and is a top priority target for antibiotic development. *Cryo*-EM structures of the *A. baumannii* F_1_F_o_-ATP synthase in three conformational states reveal unique features, which represent attractive sites for the development of novel therapeutics.

**One sentence summary:** Structure of Acinetobacter baumannii ATP synthase

## Article

*Acinetobacter baumannii* is an ESKAPE pathogen^1^ which causes a variety of hospital-acquired infections and severe treatment complications, including SARS-CoV-2 associated pneumonia^2^. A dramatic increase in multi-drug resistance (MDR) in the last decade has placed *A. baumannii* at the top of the WHO priority pathogen list, urgently requiring the development of new drugs with alternative cellular targets^3^. Recent successes in treating MDR bacteria have included novel inhibitors of the F_1_F_o_-ATP synthase, with bedaquiline (BDQ) having been approved in 2012 as the only novel last-line antibiotic for MDR *Mycobacterium tuberculosis* since rifampicin in 1971^4^. ATP synthase is highly upregulated in MDR patient strains of *A. baumannii* and, unlike many traditional targets, the enzyme is conditionally essential for survival in biofilms and dormant states^5-8^.

F_1_F_o_-ATP synthases are large, membrane-embedded macromolecular complexes that harness energy from the proton-motive force (*pmf*) and use it to synthesise ATP via a unique rotary mechanism. The simplest forms of ATP synthase are found in bacteria and plant chloroplasts and possess an F_1_ head containing subunits α_3_β_3_γδε and an F_o_ motor containing subunits ab_2_c_9-15_. Protons travel through the membrane-embedded a-subunit turning the c-ring. The rotating c-ring transfers torque to a central shaft comprising the ε- and γ-subunits which protrude into the α_3_β_3_ hexamer, inducing conformational changes in the F_1_ nucleotide binding pockets thus converting the rotary force into ATP synthesis.

In order to identify unique vulnerabilities of the *A. baumannii* ATP synthase, the complex was purified via affinity-tag and reconstituted into peptidiscs (**Figure S1A**). ATPase hydrolytic activity assays indicated that the complex was in an ‘autoinhibited state’^9^ (**Figure S1B**) The sample was analysed by single-particle *cryo*-EM (Figure S1C); 11,490 movies yielded 349,160 particles which were refined into 3 separate states, distinguished by the position of the central stalk and reached overall resolutions of 3.1 Å, 4.6 Å and 4.3 Å, respectively (**Figure S2**). Further masked refinements improved the resolution in the F_o_ region of state 1 to 3.7 Å and permitted *de novo* building of the complex into a composite map. State 1 was then used as a reference to build structures of the remaining two states (**Figure S3**).

The *A. baumannii* ATP synthase has a-subunit stoichiometry of α_3_β_3_γδεab_2_c_10_; its overall architecture resembles that of other bacterial and chloroplast ATP synthases^10-12^ (**Figure 1A**). The central stalk is rotated by almost exactly 120° between each of the three conformational states (**Figure S4A**), as seen previously^10^, suggesting common stable low-energy intermediate states in bacterial ATP synthases.

**Figure 1.**
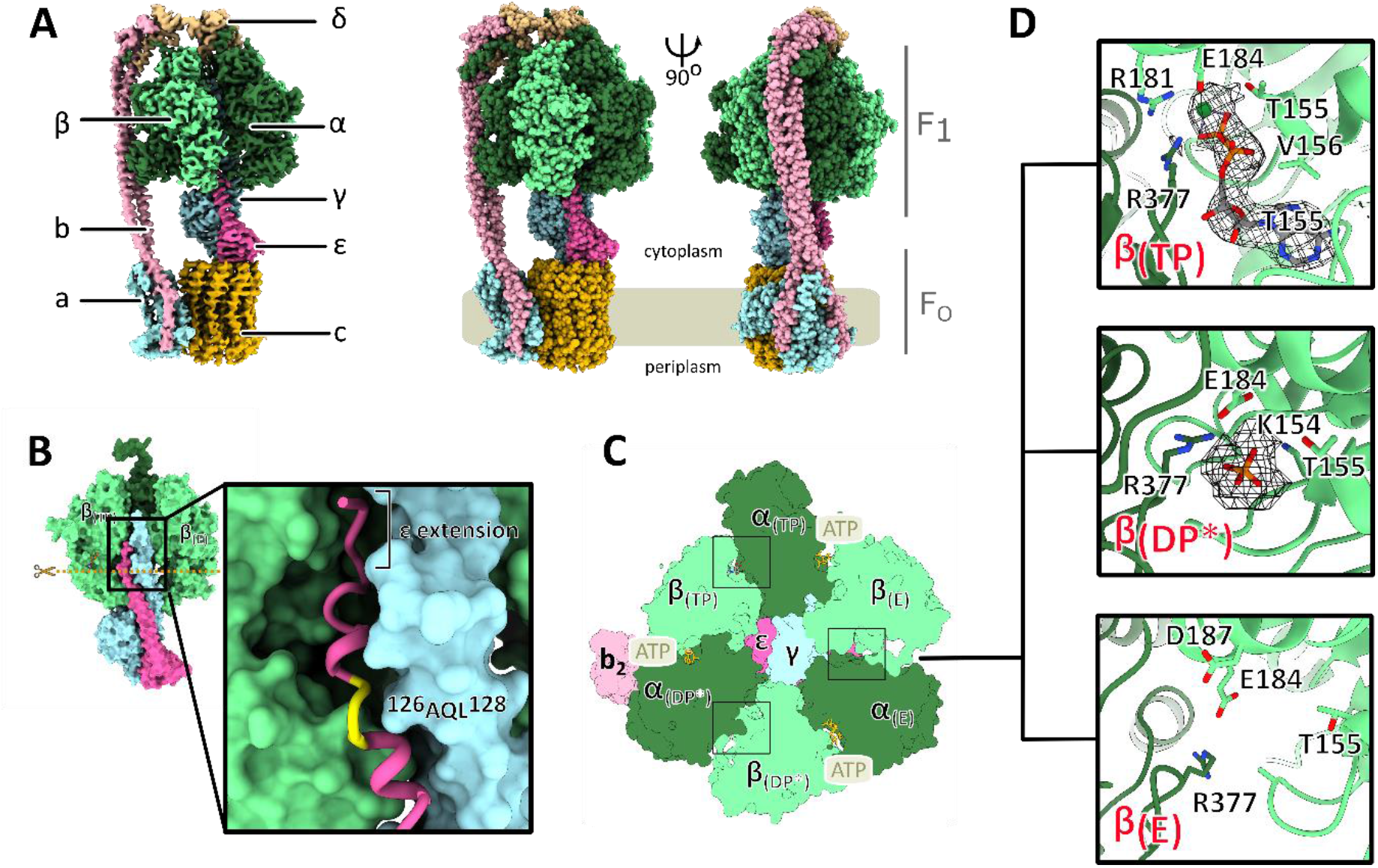
Structure of the F_1_F_o_-ATP synthase from *A. baumannii*. **A**: *Cryo*-EM map of *A. baumannii* ATP synthase coloured by subunit as indicated (left). Surface representation of the corresponding atomic structure shown as spheres in orthogonal views in corresponding colours (right). The membrane is indicated by a light brown bar. **B**: Structure of the F_1_ head including the central stalk; subunits α_DP_, β_DP_ and α_E_ removed to reveal how subunits γ (cyan) and ε (pink) insert into the head. Close-up of the C-terminal extension of ε inserting between β_TP_ and α_DP*_ (not shown) showing the unwound turn of ε ^126^AQL^128^ (yellow). **C**: Horizontal section of F_1_ as indicated by dashed line in (B) including location of nucleotide binding sites. **D**: The catalytic nucleotide binding sites with EM map features corresponding to nucleotides and phosphates shown as mesh.

In F_1_, we observe the canonical ‘Walker-Boyer’ state geometries in both the ‘*open*’ conformation (β_E_, ‘*empty*’) and the ‘*loose*’ conformation (β_TP_, with bound Mg-ADP)^13,14^ (Figure 1C). As in *Bacillus* PS3^10^, the third β-subunit (‘*tight*’ or β_DP_) is in a ‘*forced open*’ conformation (denoted as β_DP*_) and we find only weak map features in this region, which we interpret as a loosely bound inorganic phosphate ion (**Figure 1D**). This conformation is induced by an interaction between the ε-subunit and a predominantly hydrophobic loop-helix region in the β_DP_-subunit (^377^DIIAILGMDE^386^) which overlaps with the highly conserved ^385^DELAEED^391^ motif (**Figure S5**). The ‘*forced open*’ conformation of the β_DP_ site is more similar to the *open* β_E_ conformation (all-atom RMSD of 1.5 Å) than to the *loose* β_TP_ (all-atom RMSD β_DP_ to β_TP_ 2.6 Å) (**Figure S5**) but is less ‘open’ than the equivalent site in PS3 (all-atom RMSD *A. baumannii* β_DP*_ to *Bacillus* PS3 β_E_ = 0.65 Å).

The *A. baumannii* ε-subunit is similar to those seen in other bacteria, containing an extended C-terminal α-helix. However, where PS3 ε terminates, both *A. baumannii* and *Escherichia coli* unwind in a short ^126^AQL^128^ motif (**Figure 1B**)^9,10^. While the *E. coli* subunit ε bends here and protrudes ‘horizontally’ between the α_DP_- and β_TP_-subunits, *A. baumannii* ε continues further upwards into the F_1_ head, forming two more helical turns followed by a 5-residue extension (**Figure 1B**), which forms several additional interactions with the γ-, β_TP_- and the α_DP_-subunits. This may further stabilise subunit ε in the inhibitory ‘up’ position, which blocks rotation in the hydrolysis direction, while still enabling ATP synthesis^10^ (**Figure S4B**). This ‘ratchet’ mechanism is similar to that observed in PS3, but distinct from that in the mycobacterial ATP synthase, which instead relies of the formation of a temporary β-strand interaction between α and ε subunits^15^. The stronger blockage may help avoid wasteful ATP hydrolysis, enhancing *in vivo* host persistence.

The γε complex is linked to the F_o_ motor which comprises 10 copies of the c-subunit (c_10_ ring) and one copy of the proton-conducting a-subunit. Its core fold is highly conserved and resembles other ATP synthase structures, while the proton conducting channels are most similar to those in other bacterial a-subunits (**Figure 2A**). Bacterial ATP synthases contain sequence insertions in the N-terminus of the a-subunit and *A. baumannii* has the largest insertion of the structurally characterised ATP synthases (**Figure S6A**), resulting in a repositioning of the first short α-helix towards the periplasm (**Figure 2B**). Importantly, this shift relocates the entry site of the periplasmic proton channel: in the mammalian ATP synthase protons enter from behind this mini-helix, whereas in *A. baumannii* they approach from the front, as viewed from the c-ring (**Figure 2B**). Positioning of this mini-helix also appears important in determining the channel entry point in other ATP synthases (**Figure S7**). As several inhibitors of ATP synthases bind to the proton entry channel, including organotin compounds currently employed as pesticides^16,17^, the identification of a distinct entrance may provide an avenue to design specific inhibitors which target bacterial ATP synthases but not those of the mammalian host (**Figure 2B, Figure S7**).

**Figure 2.**
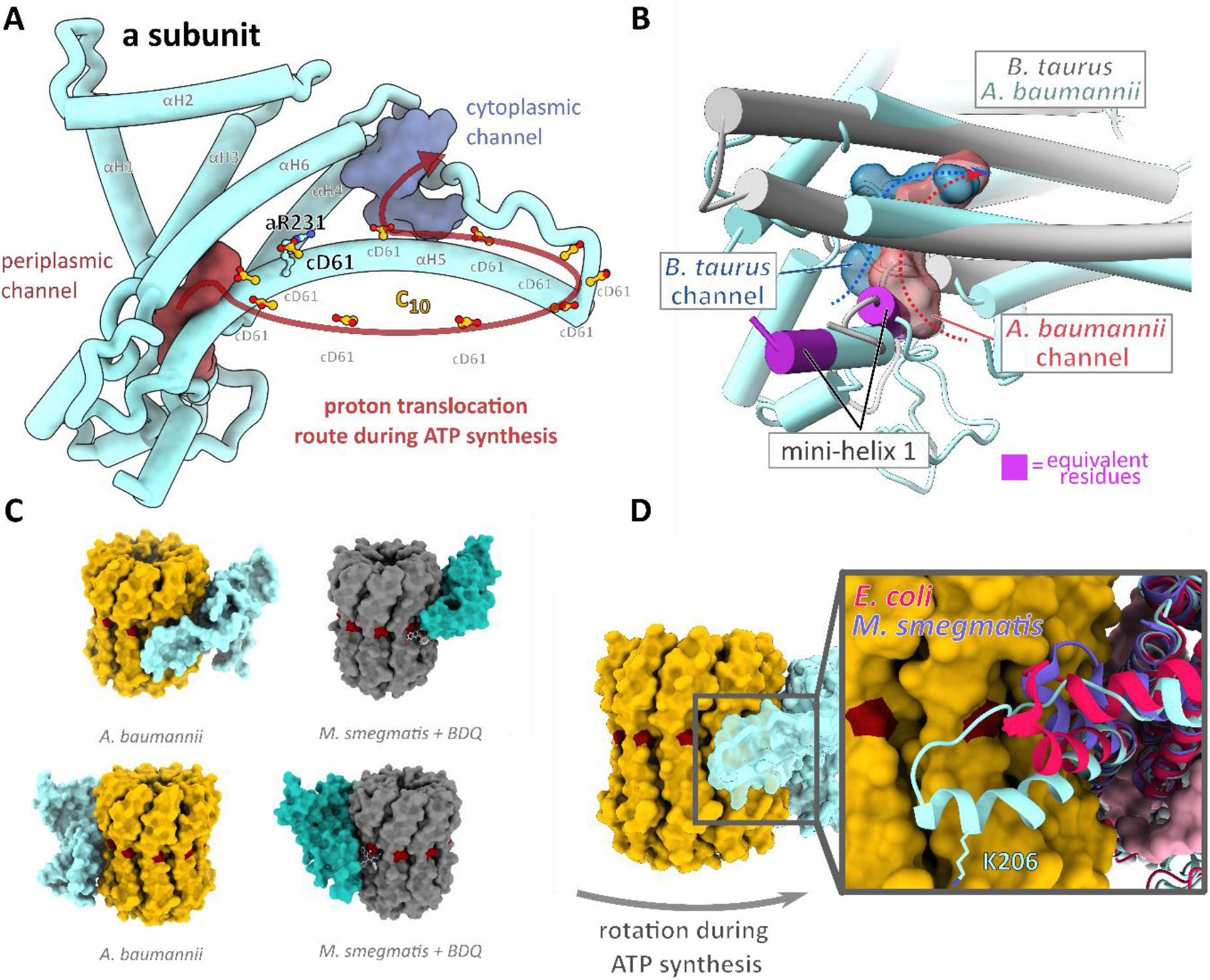
Structure of F_o_ motor and *Acinetobacter* specific features of the entry/exit channel sites. **A**: Ribbon representation of the a-subunit with the key conserved aR231 and the protonatable cD61 carboxylates in the c_10_ ring shown in ball and stick representation. Periplasmic and cytoplasmic channels, are shown in maroon and purple, respectively. Arrow indicates direction of rotation during ATP synthesis. **B**: Surface rendering of the periplasmic channels in the a-subunits of *B. taurus* (dark blue) and *A. baumannii* (maroon) indicating alternate exit pathways. Equivalent residues forming the first mini-helix are coloured in purple. **C**: Comparison of a/c_10_ interface in *A. baumannii* (yellow and light blue) and in *M. smegmatis* (dark grey and turquoise, PDB:7JGA) showing binding site of bedaquiline (BDQ) in *M. smegmatis* (light grey ball and sticks), proton carrying residues shown in dark red. Bottom panels are rotated by 180° around the membrane normal **D**: Surface representation of c-ring (yellow) and a-subunit (translucent cyan) viewed from within the membrane plane. *Insert* – structural alignment of a-subunits from *A. baumannii, E. coli* (PDB:6OQR) and *M. smegmatis* (PDB:7JGA) indicate extent of a-subunit loop insertion. K206 shown in stick representation.

The *A. baumannii* a-subunit contains an additional loop extension between aH5 and aH6 (**Figure 2C and D**). The loop extension is formed by predominantly hydrophobic residues plus a charged lysine residue (^200^PSNPVAKALLIP^211^) and is conserved in the *Acinetobacter* genus and in the *Moraxellaceae* family but is fully or partially absent in other ATP synthases (**Figure S6A**). A structural alignment of a-subunits from several bacterial ATP synthases (**Figure 2D**) reveals that the *A. baumannii* loop is significantly longer than in other bacteria and may even reach the periplasmic membrane edge (**Figure S8**). In cells, such proximity may provide privileged access to small molecules and biologics, which is absent in other species. Additionally, K206 appears to stretch towards the bilayer leaflet, suggesting it might even interact with phospholipid headgroups (**Figure 2D**).

The success of Bedaquiline (BDQ) in treating tuberculosis has ignited interest in targeting ATP synthases in ESKAPE pathogens. Unlike most ATP synthase inhibitors^17^, BDQ binds with high specificity to mycobacterial ATP synthases. In structures of BDQ bound to the *M. smegmatis* ATP synthase, the highest affinity binding sites were found at the interface between c_10_ and the a-subunit (Figure 2C)^15^. Furthermore, the broad-spectrum antimalarial antibiotic mefloquine and the new antibiotic lead tomatidine^18^ are suggested to act in a similar manner, targeting the a/c-ring interface^19^.

As the a-subunit loop insertion is conserved within the *Acinetobacter* family but is absent or structurally diverse in other bacteria and the mitochondrial ATP synthase, the unique a/c_10_ interface in *A. baumannii* represents a prime target for the development of highly specific inhibitors. BDQ belongs to a class of small molecules known as diarylquinolines (DARQs). A targeted screen of ∼700 DARQs yielded compounds that specifically inhibited the ATP synthase from *S. aureus* and *Streptococcus pneumoniae* but showed minimal activity towards *M. tuberculosis*^20^. These data demonstrate that inhibitors which bind at the c-ring or the a/c_10_ interface can be optimised to target specific ATP synthases beyond *Mycobacteria* and future structure-based drug design may facilitate the development of compounds targeting this interface in ESKAPE pathogens^1^. The structure of the *A. baumannii* ATP synthase reveals novel and unique features of a bacterial ATP synthase and will inform future efforts to develop much needed antibiotics to treat MDR *Acinetobacter*.

## Acknowledgments

This work is dedicated to our late collaborator Éric Marsault (Sherbrooke University, Canada). We thank Paul Simpson and the Centre for Structural Biology (CSB) electron microscopy facility at Imperial College London for EM support. We are also grateful to Nora Cronin and the London Consortium for electron microscopy (LonCem) at the Crick Institute (London, UK), and the Electron Bio-Imaging Centre (eBIC) at Diamond Light Source (Harwell, UK) for their continued support and microscopy time. Suhail Islam is acknowledged for IT support. Nita Shah and Anaïs Menni are thanked for processing advice to JD. This work was funded by a Wellcome UK Investigator grant to TM [WT110068/Z/15/Z].

## Author contributions

JKD, OLU and LPA grew cells, purified and optimised the protein sample. JKD collected and analysed data, built and analysed the structure and provided a manuscript draft. BPP analysed the structure, made figures, and wrote the final paper version. OLU performed biochemical assays. LPA and AF provided constructs, helped with *Acinetobacter* genetics, provided method sections and comments for the manuscript. MB built, refined and analysed the structure and contributed to the manuscript. TM obtained funding, analysed and interpreted data, wrote the paper and directed the project.

## Abbreviations

ATP: Adenosine-5’-triphosphate
*cryo*-EM: electron *cryo*-microscopy
MDR: multi drug resistance
(XDR): extensively drug resistance

