## Supplemental Materials for "Structure of ATP synthase from ESKAPE pathogen *Acinetobacter baumannii*"

[1] *Department of Life Sciences*
*Imperial College London*
*Exhibition Road*
*London SW7 2AZ, United Kingdom*

[2]*MRC Centre for Molecular Bacteriology and Infection*
*Department of Life Sciences*
*Imperial College London*
*London SW7 2AZ, United Kingdom*

[3]*National Heart and Lung Institute*
*Imperial College, London, UK*

[4]*Department of Biochemistry*
*University of Oxford*
*South Parks Road*
*Oxford OX1 3QU, United Kingdom*

[5]*Private University in the Principality of Liechtenstein*
*Triesen, Liechtenstein*

*These authors contributed equally.

[Ŧ]corresponding author:
Thomas Meier


**Supplementary Figures:**

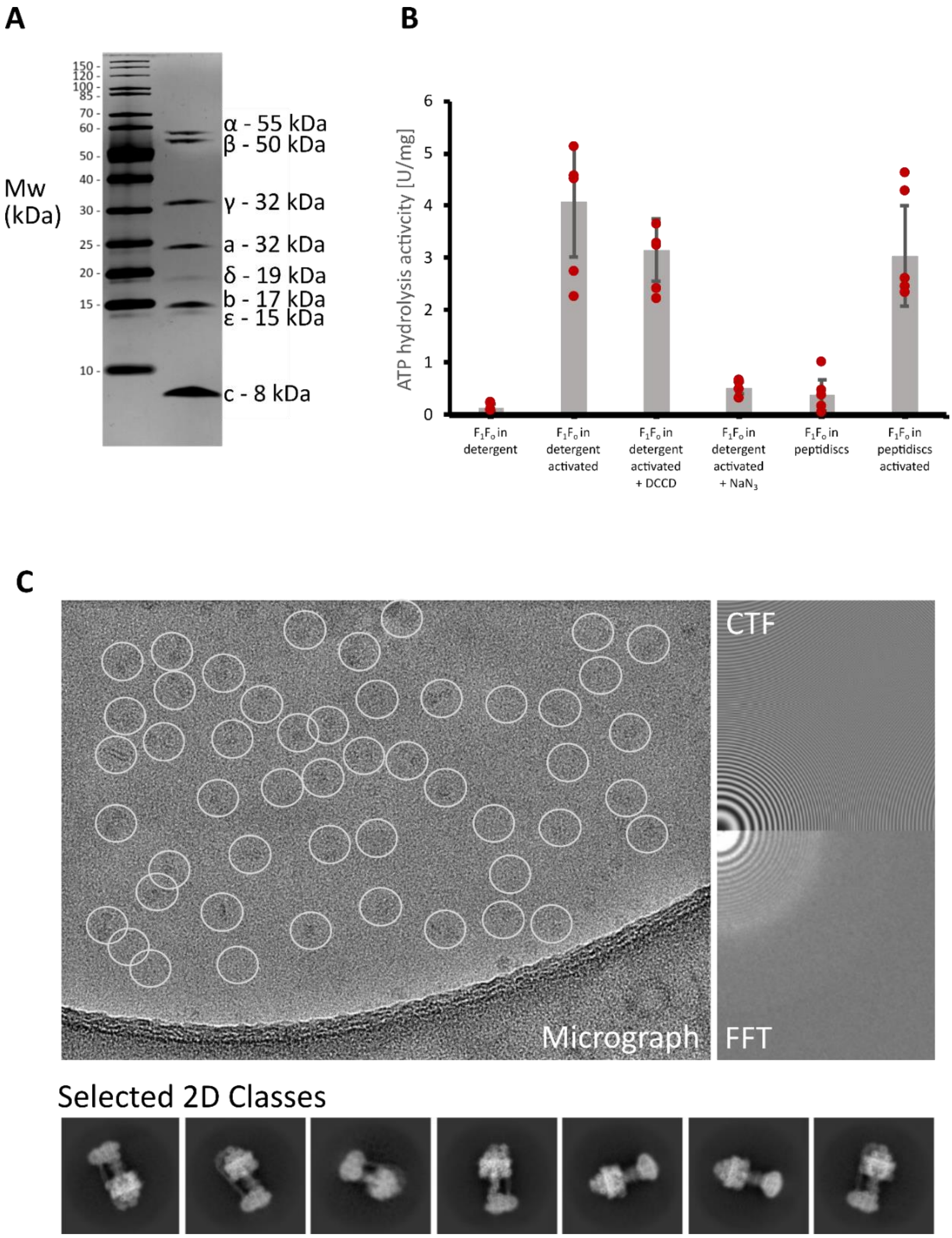

**Figure S1 – Purification, biochemical characterisation, and sample quality of the *A. baumannii* ATP synthase. A:** Detergent-solubilised ATP synthase on SDS-PAGE (stained with silver). **B:** ATPase hydrolytic activity in the presence of activators (detergent) and inhibitors (DCCD and $NaN_3$). The ATPase enzyme activity of the purified complex was blocked both in 4-trans-(4-trans-propylcyclohexyl)-cyclohexyl α-maltoside (tPCC-α-M) and peptidiscs but could be restored by addition of lauryl-dimethyl-amine oxide (LDAO) and trypsin to a specific activity of ~4 U/mg in tPCC-α-M and ~3 U/mg in peptidiscs (labelled activated on x-axis). In this activated form, the sample could not be inhibited by the $F_o$ c-ring inhibitor *N*,*N*'-dicyclohexylcarbodiimide (DCCD) but still showed full inhibition by $NaN_3$, which binds the β-subunit nucleotide binding sites, indicating that $F_1$ is uncoupled from $F_o$ upon enzyme activation by LDAO/trypsin. **C:** Representative micrograph showing picked particles (white circles), CTF estimation, FFT of raw image and selected 2D classes of *A. baumannii* ATP synthase processed in CryoSPARC version 2.15.

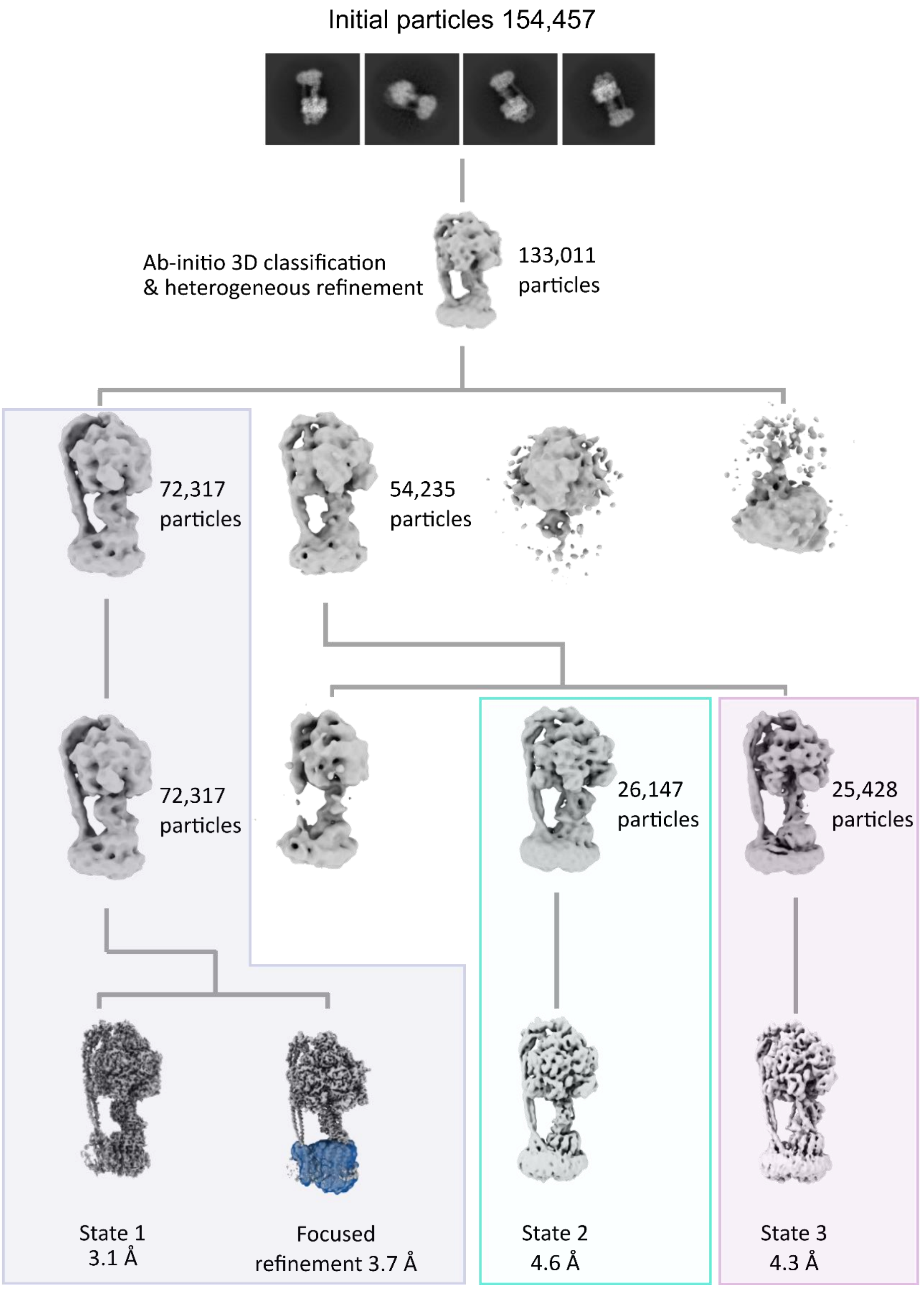

**Figure S2 – Flowchart for *cryo*-EM data processing.** Particles were subjected to multiple rounds of picking and refinement as indicated. Different rotational states were separated by 3D classification, with each class selected for further processing to high resolution as shown. Once final particles had been selected final rounds of polishing, non-uniform refinement and post-processing resulted in 3 maps with global resolution ranges of 3.1 Å – 4.6 Å. Further focused refinement of the membrane-embedded $F_O$ region in the state 1 map (blue) improved the resolution to 3.7 Å for this region. All processing was conducted in CryoSPARC version 2.15.

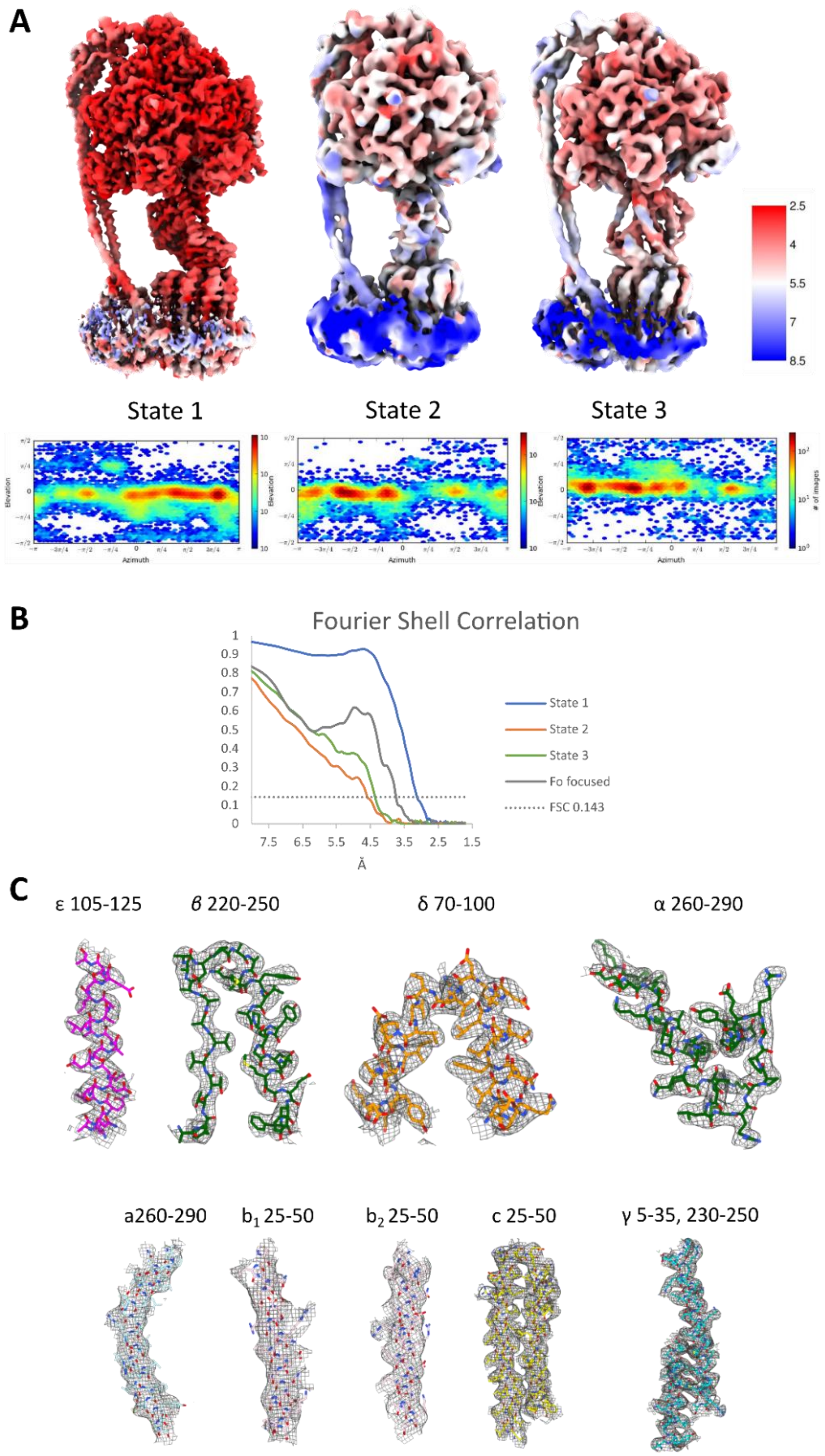

**Figure S3 – Quality of EM maps of *A. baumannii* ATP synthase. A**: Maps of 3 rotational states of ATP synthase coloured according to local resolution (red = high, blue=low). Below each map is the corresponding Euler angle plot (CryoSPARC version 2.15) indicating angular distribution of views observed in 3D reconstruction. **B**: Fourier Shell Correlation (FSC) curves for 4 EM maps as labelled. **C**: Details of map features of state 1 in different subunits with examples of atomic models built in the *cryo*-EM maps (displayed amino acid sequences are indicated).

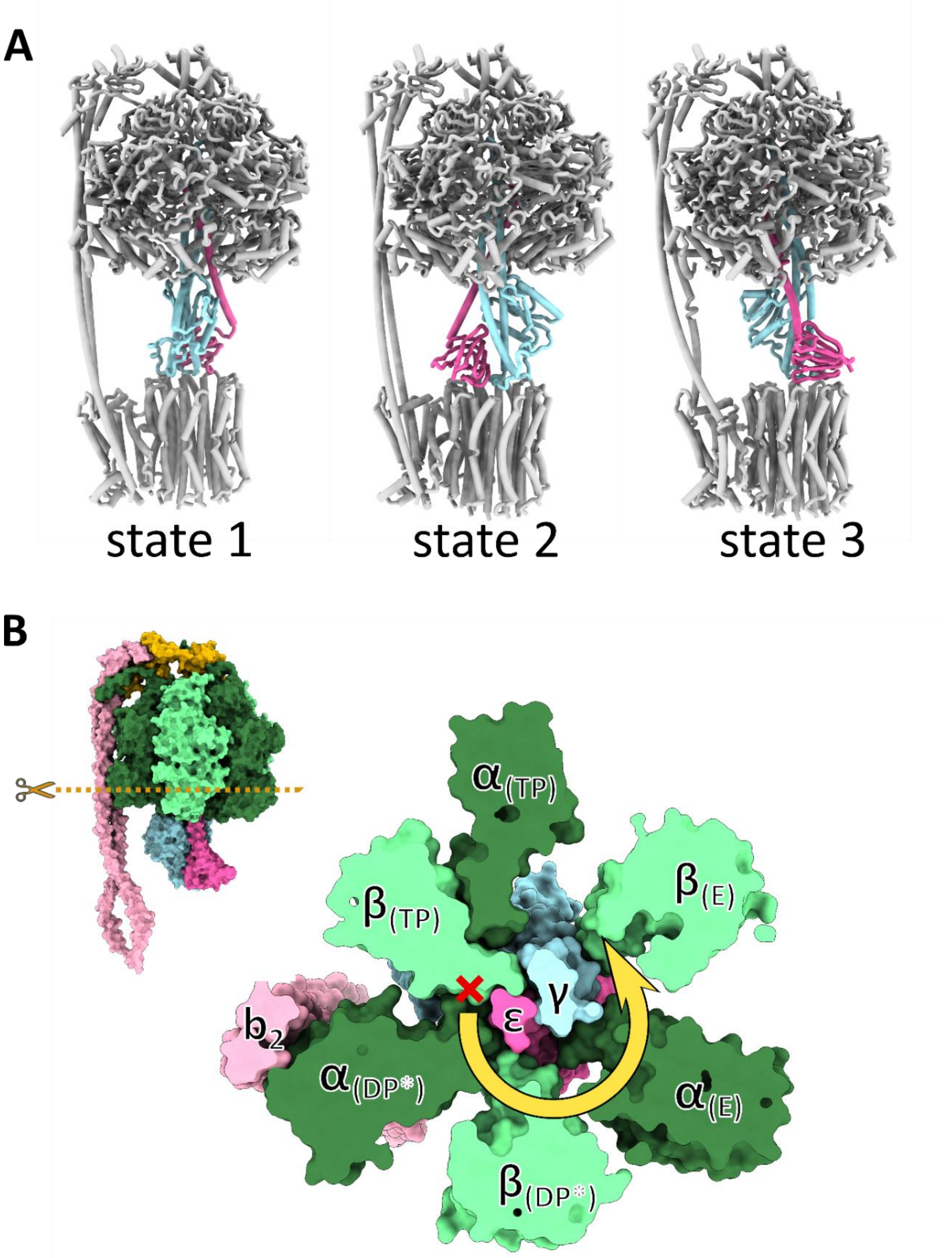

**Figure S4 – Rotational states and unidirectional inhibition of ATP synthase activity of *A. baumannii* $F_1$. A**: Cartoon representation of the 3 rotational states observed for *A. baumannii* ATP synthase, differing in the relative positions of the central stalk (shown in light blue (γ) and pink (ε)) and the conformations of the α and β subunits. Structural alignments of the γ subunit between states revealed that the three states differ by almost exactly 120˚. **B**: Horizontal section at indicated position reveals mechanism of unidirectional inhibition. Steric clashes between ε and $\beta_{TP}$ prevent rotation of the central stalk in the ATP hydrolysis direction (red cross) but permit unlocking and rotation in the ATP synthesis direction (yellow arrow).

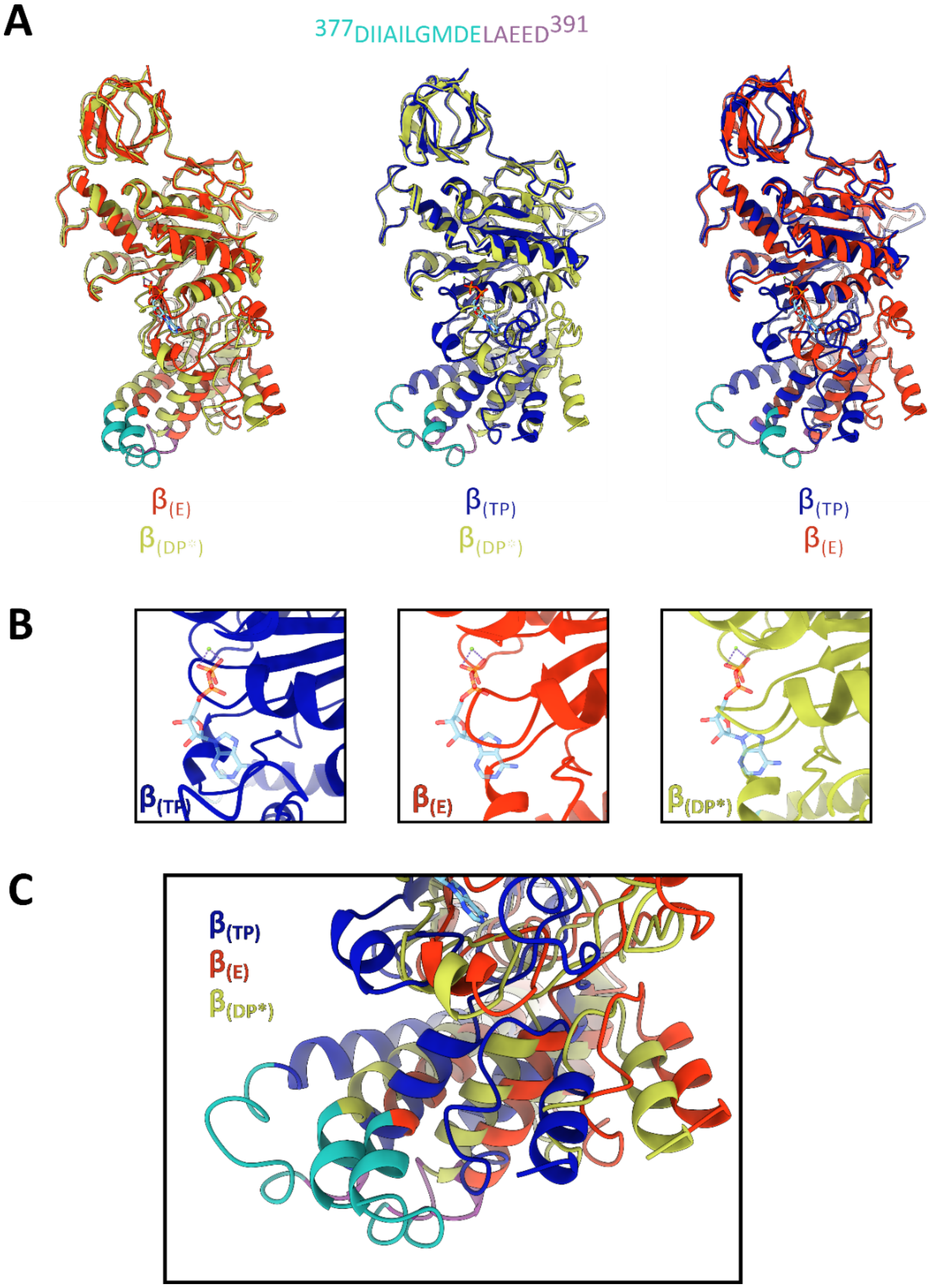

**Figure S5 – Conformational changes in the catalytic β subunits of *A. baumannii* ATP synthase. A:** Structural alignments of $β_{TP}$ (blue), $β_{DP*}$ (yellow) and $β_E$ (red) demonstrate conformational transitions during the ATP synthase catalytic cycle. ADP is shown in stick representation in its binding location in $β_{TP}$ in all 3 alignments for reference only. **B**: Close up of nucleotide binding pockets for all 3 states with ADP as observed in $β_{TP}$ shown in cyan (nitrogen = dark blue, oxygen = red, phosphorous = orange). Steric clashes between the peptide backbone and the ADP in $β_E$ and $β_{DP*}$ clearly indicate incompatibility with ATP binding. **C**: Close-up of conformational changes in the ε-proximal helix-turn-helix motif in the β-subunit ($^{377}$DIIAILGMDE$^{386}$ – shown in cyan) which prevent bidirectional rotation of the central stalk when in the $β_{TP}$ conformation, but not in the $β_E$ or $β_{DP*}$ conformations. The state 1 structure was used to generate this figure. All alignments were performed in UCSF ChimeraX.

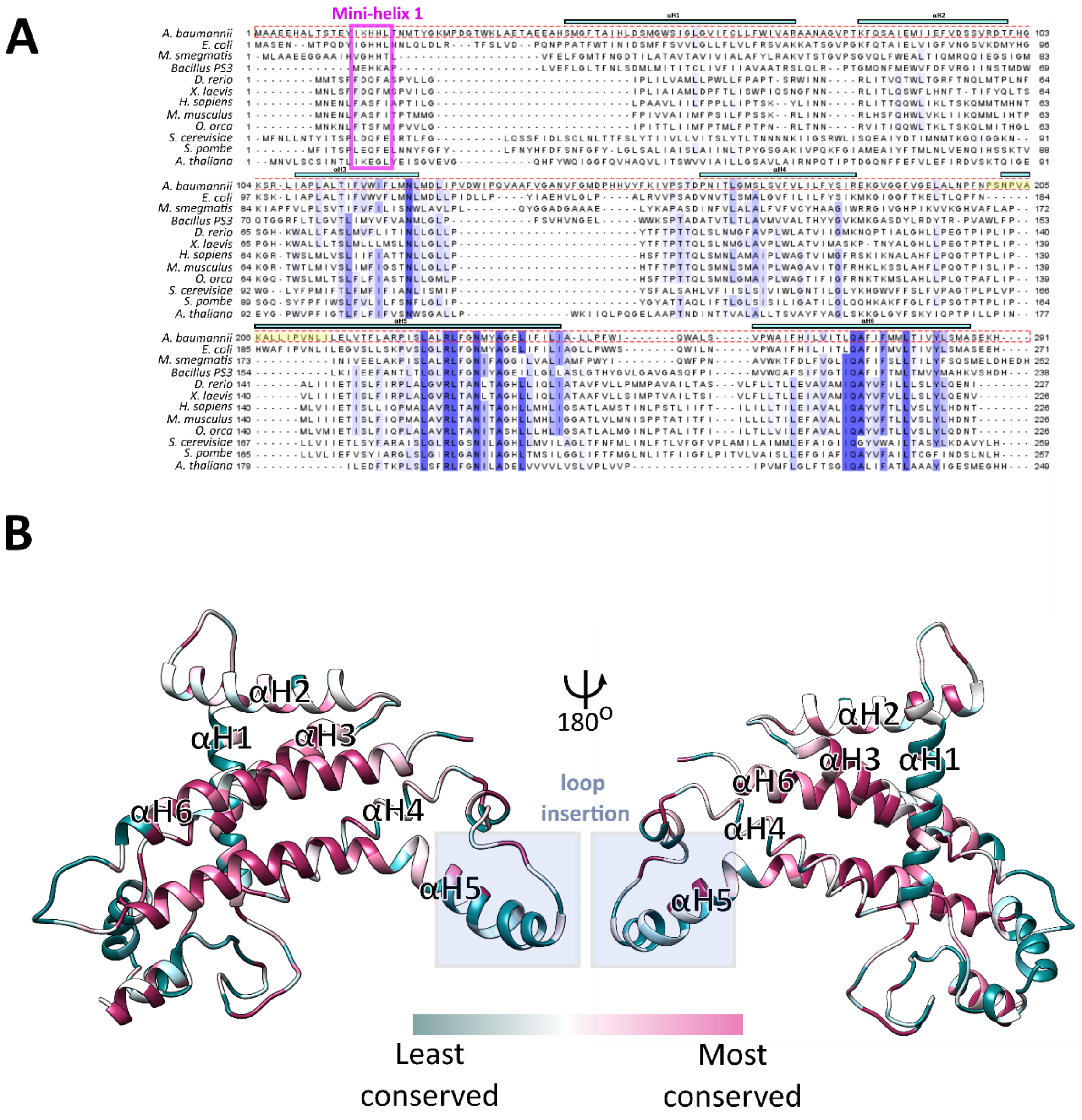

**Figure S6 – Amino acid alignment and residue conservation analysis of subunit a of the *A. baumannii* ATP synthase. A:** Multiple sequence alignment (produced using ClustalX2 and Jalview) of the a-subunit of the ATP synthase from model organisms with residues coloured by percentage identity across species (white = non conserved, blue = conserved, red dashed box = *A. baumannii* sequence). Pink box indicates residues coloured purple in **Fig. 2B** which form mini-helix 1 in *A. baumannii* and *B. taurus.* The locations of the a-subunit helices (aH1 to aH6) are indicated above the alignment. Yellow highlight indicates *A. baumannii* loop insertion between aH5 and aH6. **B**: Cartoon representation of the *A. baumannii* a-subunit coloured according to conservation (bottom) from least conserved (turquoise) to most conserved (pink), generated using the CONSURF server[1]. The core parts of helices aH5 and aH6 are the highest conserved regions in the a-subunit. Blue box indicates location of a-subunit loop insertion.

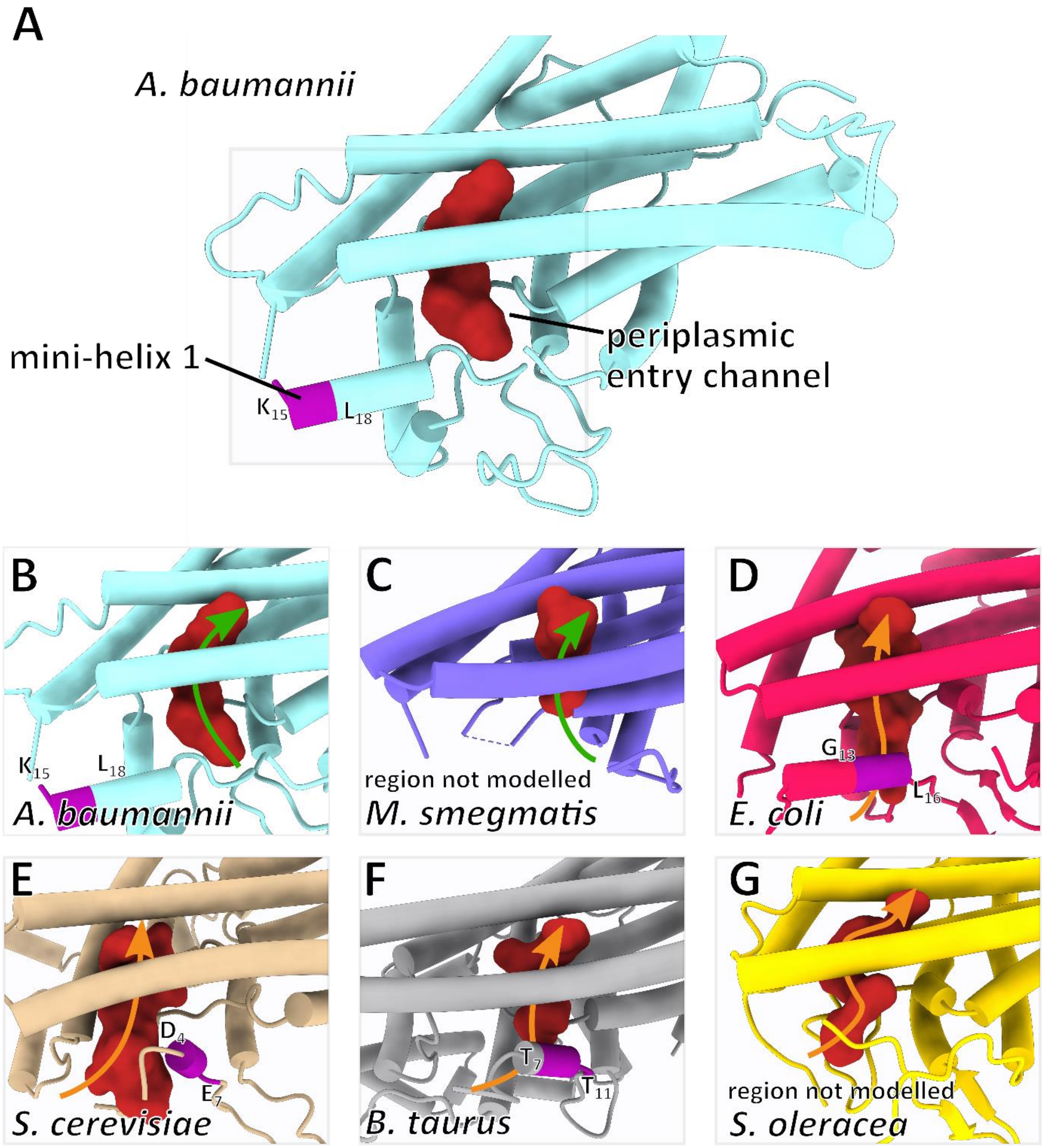

**Figure S7 – Position of a-subunit mini-helix 1 guides location of periplasmic entry channel. A:** Periplasmic entry channels of a-subunits of ATP synthases with corresponding residues, which form a-subunit mini-helix 1, are coloured in purple and surface rendering of channels are coloured brick red. All subunits are viewed from the perspective of the c-ring and are coloured as in Figure 2D. **B-G:** Close up views of channels from ATP synthase a subunits. 'Front' entrance channels are indicated by a green arrow, 'back' entrance channels are indicated by an orange arrow. In general cases, where mini-helix 1 is shifted towards the periplasm as a result of an N-terminal sequence insertion (*A. baumannii* – PDB: 7P2Y)(**B**) and/or missing in the structure (*M. smegmatis* – PDB: 7JGA)(**C**) the periplasmic channel appears to emerge on the right (in front of the helix position). In the *E. coli* (PDB: 6OQR) channel (**D**) as well as the mitochondrial ATP synthases of yeast (*S. cerevisiae* – PDB: 6B9H)(**E**) and bovine (*B. taurus* – PDB: 6ZQM)(**F**), a shift of mini-helix 1 back into the core fold creates an obstruction, which guides the channel away towards the left (behind the helix). Finally, although this position of mini-helix 1 does appear to guide the channel towards the left, formation of a helix in this location is not strictly necessary as the spinach chloroplast ATP synthase structure (*S. oleracea* – PDB: 6FKF)(**G**) indicates that a different region of the backbone can still occlude the right facing channel without forming the mini-helix motif.

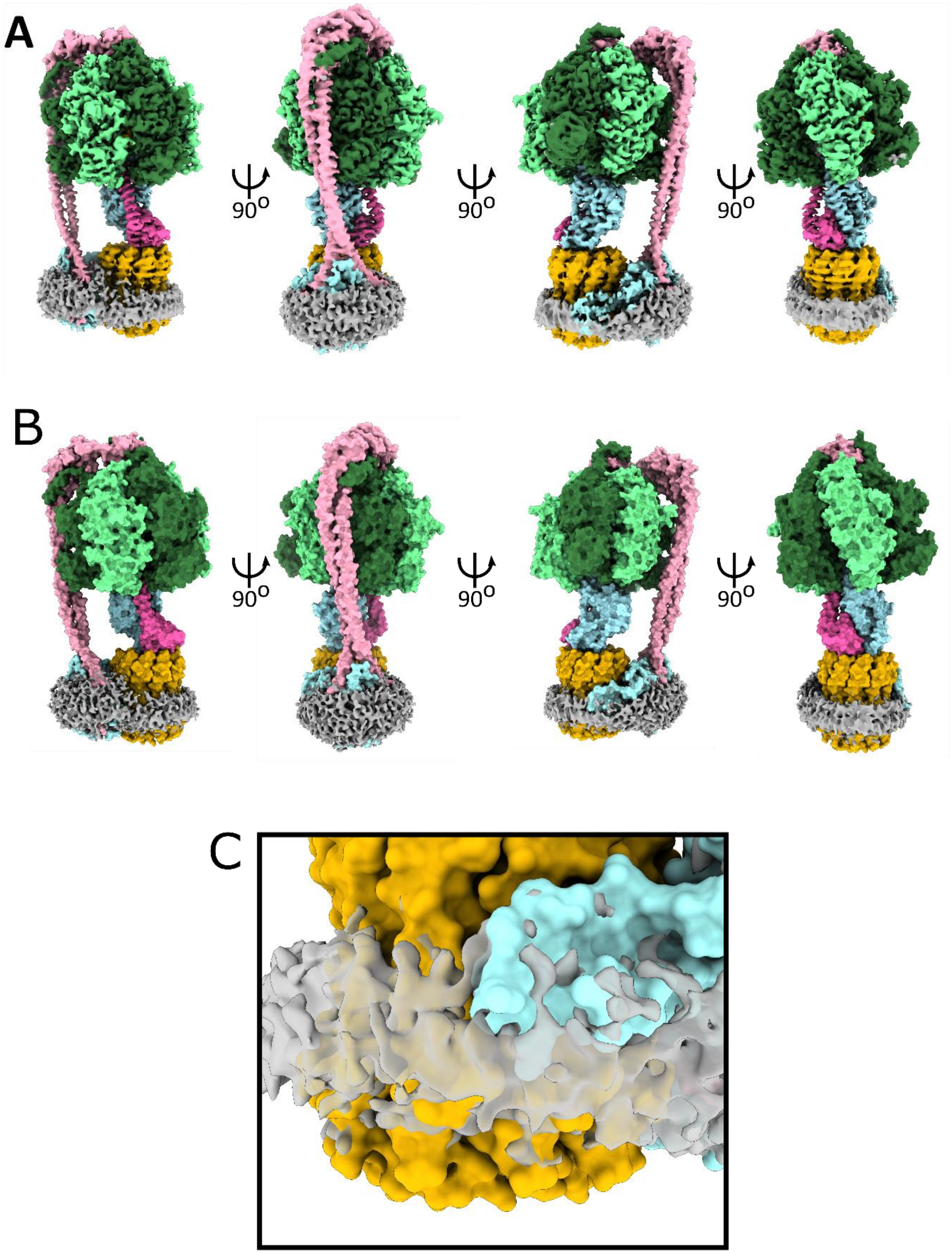

**Figure S8 – Peptidisc boundaries of the *A. baumannii* ATP synthase**. **A**: Rotated side views of the *cryo*-EM map for *A. baumannii* in peptidisc (state 1), coloured by proximity to main chain in same colour scheme as **Figure 1A**. Map was coloured using UCSF ChimeraX with a 'zone' radius of 5 Å. Note that the peptidisc map is more than 5 Å away from the peptide backbone apart from at the a-subunit loop extension near the periplasmic membrane envelope (view third in from the left). **B**: Surface representation of the *A. baumannii* structure with *cryo*-EM map for the peptidisc superimposed. **C**: Close up view of a surface representation (as seen in panel B, third position) of the a-subunit loop insertion superimposed with the peptidisc *cryo*-EM map.

**Table S1 - *Cryo*-EM model building and statistics**

| | State 1 | State 2 | State 3 |
|---|---|---|---|
| **Data collection** | | | |
| Accession numbers | 7P2Y<br>EMD-13174 | 7P3N<br>EMD-13181 | 7P3W<br>EMD-13186 |
| Electron microscope | Titan Krios G3 | Titan Krios G3 | Titan Krios G3 |
| Electron detector | Gatan K3 | Gatan K3 | Gatan K3 |
| Nominal Magnification | 85,000 | 85,000 | 85,000 |
| Voltage (kV) | 300 | 300 | 300 |
| Defocus range (µm) | -1.5 - 3 | -1.5 - 3 | -1.5 - 3 |
| Pixel size (Å) | 0.85 | 0.85 | 0.85 |
| Total Dose ($e^-$/ Å²) | 60 | 60 | 60 |
| Symmetry imposed | C1 | C1 | C1 |
| Total micrographs | 11,490 | 11,490 | 11,490 |
| Initial particle images (no.) | 349,160 | 349,160 | 349,160 |
| Final particle images (no.) | 72,317 | 26,147 | 25,428 |
| Map resolution (Å) | 3.1 | 4.6 | 4.3 |
| FSC threshold | | 0.143 | |
| Map resolution range (Å) | 2.6-8.5 | 3.9-8.5 | 3.5-8.5 |
| Map sharpening *B* factor (Å$^2$) | -60 | -102 | -74 |
| **Image processing** | | | |
| Initial model | *De novo* | *De novo* | *De novo* |
| Model composition | | | |
| Protein chains | 22 | 22 | 22 |
| Non hydrogen atoms | 37183 | 37126 | 37161 |
| Protein residues | 4904 | 4904 | 4904 |
| Ligands | 3 Mg-ATP<br>1 Mg-ADP<br>1 $P_i$<br>3 $H_2O$ | 3 Mg-ATP<br>1 Mg-ADP<br>1 $P_i$<br>3 $H_2O$ | 3 Mg-ATP<br>1 Mg-ADP<br>1 $P_i$<br>3 $H_2O$ |
| R.m.s. deviations | | | |
| Bond lengths (Å) | 0.004 | 0.005 | 0.005 |
| Bond angles (°) | 0.716 | 0.861 | 0.838 |
| Ramachandran plot | | | |
| Favoured (%) | 96.2 | 92.5 | 93.7 |
| Disallowed (%) | 0 | 0.06 | 0.02 |
| Validation | | | |
| Rotamer outliers (%) | 0.08 | 0.21 | 0.13 |
| MolProbity Clashscore | 10.37 | 30.55 | 26.71 |

### Materials and methods

#### Chemicals, enzymes and strains

If not stated otherwise all chemicals were purchased from Sigma Aldrich and Life Technologies / Thermo Fisher Scientific.

#### Bacterial strains and growth conditions

*Acinetobacter baumannii* ATCC 17978 was grown in tryptone soy broth (TSB), Terrific Broth (TB) or Lysogeny Broth (LB) agar plates at 37°C. *Escherichia coli* Top10 was grown in LB at 37°C with the addition of agar as required. Media were supplemented with antibiotics tetracycline 15 µg/ml or kanamycin 50 µg/ml as required.

#### Molecular Cloning

*A. baumannii* genomic DNA isolation was performed using the PureLink Genomic DNA mini kit (Life Technologies). Isolation of plasmid DNA was carried out using the QIAprep spin miniprep kit (Qiagen). The β-subunit encoding gene of *A. baumannii* ATP synthase (3815454- 3814060, GenBank: CP053098.1) was amplified with KOD Hot Start DNA Polymerase (Novagen) as described by the manufacturer with the inclusion of 1 M betaine using the BetaFstrep forward primer 5'-TCTGACTCGAGTAACAGGAGGAATTAACCATGTGGAGCCACCCGCAGTTCGAAAAGAGTAGCGGTCGTATC-ATTC-3' and the BetaR reverse primer 5'-CAGATCTGCAGTTAGAGTTTCTCAGCTTTAGCAA-3'. These primers introduce the optimised pBAD 18 bp Ribosome Binding Site (RBS) and then the sequence encoding a StrepII tag at the gene 5' end of the gene. The amplified gene was inserted into pCR-Blunt II-TOPO and transformed into *E. coli* Top10 and confirmed by colony PCR screening with standard *Taq* polymerase (NEB) with the addition of 2% dimethyl sulfoxide (DMSO). The correct insert was then digested from purified plasmid using *Xho*I and *Pst*I restriction endonucleases according to the manufacturer's specifications (Roche). The insert was then ligated into digested pBBR-MCS3[2] prior to transformation into Top10. The confirmed sequence of plasmid pBBR_AbATPbetaSII was then introduced into *A. baumannii* cells *via* electroporation. DNA sequences were confirmed by sequencing of all polymerase amplified regions (GATC Biotech).

#### Isolation of ATP synthase from *Acinetobacter baumannii*

Cultures of *A. baumannii* cells containing vector pBBR_AbATPbetaSII were grown in TB media containing tetracycline 15 µg/ml overnight and harvested by centrifugation at 5,000 x g. All subsequent steps were performed at 4°C. 5 g wet cells were resuspended in buffer A (50 mM 4-(2-hydroxyethyl)-1-piperazineethanesulfonic acid (HEPES) buffer pH 6.8, 150 mM KCl, 5 mM $MgCl_2$) supplemented with a spatula tip of DNase I and cOmplete™ Protease Inhibitor Cocktail (Roche). Cells were lysed by passaging three times through a French Pressure Cell at 20,000 psi and centrifuged at 10,000 x g for 1 h. The remaining supernatant was first filtered through 0.2 µm filters and then centrifuged at 200,000 x g for 1 h to harvest membranes. The supernatant was discarded and membranes were resuspended and solubilised in buffer A, which was supplemented by 1% (w/v) of trans-4-(trans-4'-propylcyclohexyl)cyclohexyl-α-*D*-maltoside (tPCC-α-M, Glycon, Luckenwalde, Germany) by gentle agitation for 60 min. The unsolubilised membranes were removed by ultra-centrifugation (200,000 x g, 30 min). The supernatant was next loaded onto a Strep affinity column (IBA Lifesciences) equilibrated in buffer B (50 mM HEPES pH 6.8, 100 mM KCl, 5 mM $MgCl_2$ with 0.05% (w/v) tPCC-α-M) and eluted with buffer B containing 2 mM desthiobiotin using an Äkta Pure chromatography system (GE Healthcare). The eluate was loaded onto a MonoQ 5/50 GL column (GE Healthcare), which was previously equilibrated with buffer B. The product was then gradually eluted with buffer B plus 1 M KCl. Fractions containing the ATP synthase were pooled and desalted using a PD-10 desalting column (GE Healthcare). The enriched ATP synthase was next bound to 0.3 ml buffer B equilibrated Q Sepharose material for 1 h. Fractions of 50 to 100 µl were eluted with buffer B plus 500 mM KCl. The purified *A. baumannii* ATP synthase sample was next reconstituted into peptidiscs

(Peptidisc Biotech)[3] using a ratio of peptidisc to sample of 2:1 (w/v). Finally, sample polishing was performed by size exclusion chromatography (Superose 6 Increase 3.2/300, buffer A, without detergent) and fractions containing the enzyme were selected by SDS-polyacrylamide gel electrophoresis (SDS-PAGE)[4] and subsequently pooled.

**Other biochemical methods**

ATP hydrolytic activity was assayed as described[5]. ATP hydrolysis activity was induced by the addition of 0.5% (w/v) lauryldimethylamine oxide (LDAO) and trypsin digestion (1:1 (w/w)) at 37°C. Digestion was stopped by the addition of aprotinin (1:5 (w/w)) after 5 minutes for 2 minutes on ice. Protein samples were analysed by SDS-PAGE[4]. The protein concentration was determined by the bicinchoninic acid (BCA) assay (Thermo Fisher/Pierce) or $A_{280nm}$ absorbance via NanoDrop™ (Thermo Fisher).

**Sample preparation & *Cryo*-EM**

Ultra-thin carbon support film, 3 nm on lacey carbon grids (Agar Scientific) were plasma cleaned in a hydrogen environment for 30 s. 4 µl of 0.05 mg/ml purified sample was pipetted on the grid in a Vitrobot III (Thermo Fisher/FEI) in a 100% humidity chamber at 8°C and frozen in liquid ethane. Images were recorded in a Titan Krios G3 microscope operated at 300 kV (Thermo Fisher/FEI) with electron-optical alignments adjusted with Sherpa (Thermo Fisher) on a Gatan K3 direct electron detector in electron counting mode at a nominal magnification of x85,000, corresponding to a calibrated pixel size of 0.425 Å in super resolution mode. 11,490 dose fractionated movies were recorded using EPU (Thermo Fisher/FEI) with an electron flux of 1.42 $e^-$ x $pixel^{-1}$ x $s^{-1}$ over 42 fractions corresponding to a total dose of ~60 $e^-/A^2$ in a defocus range of –1.4 to –3 µm.

**Image processing**

Image processing was performed using CryoSPARC throughout the complete image processing procedure[6]. First, beam-induced motion was corrected and dose-weighted images from movies for initial image processing were generated. Then, CTF parameters for each movie were determined. Next, particle images were automatically picked and extracted with a box size of 450 × 450 pixels. The data set was cleaned by 2D classification and several iterations of *ab-initio* reconstruction and heterologous refinement resulted in three final maps (**Figures S2 and S3**). The maps were refined using non-uniform refinement from 72,317 (*state 1*), 26,147 (*state 2*), and 25,428 (*state 3*) polished particles. Gold-standard Fourier shell correlations were calculated from two independently refined data sets to determine the overall resolution of the reconstructions according to the 0.143 FSC criterion[7]. To improve the reconstruction of the membrane region, focussed refinements using a mask excluding the $F_1$ subcomplex ($\alpha_3\beta_3\gamma\varepsilon$) and a soft-edged mask around the membrane-embedding $ab_2c_{10}$ subcomplex was applied before local realignment. Local resolution was assessed using the built-in routine and maps were sharpened in CryoSPARC. All *cryo*-EM data collection and refinement parameters are available in **Table S1**.

**Model building and refinement**

The structure was built into the EM maps in Coot[8] using a combination of Phyre2[9] and based on homologous bacterial and chloroplast structures[10-12]. Parts of the a, b, c, γ, ε subunits, δ, γ and C and N-termini of the α and β subunits were built manually *de novo*. The structure was refined by Namdinator[13] and Phenix real space refinement[14] followed by manual polishing in Coot, using a composite map of the 3.1 Å state 1 map and the 3.7 Å focused map of the $F_o$ membrane region. MolProbity[15] was used for validation. Water-accessible regions of the membrane intrinsic $F_o$ subcomplex were probed by mapping the interior surface using HOLLOW[16]. Figures and movies were made with Pymol, Chimera or ChimeraX[17-19]. Rotational analysis was performed according to[11].

**Hollow channel analysis**

All proton channel analyses were conducted using Hollow1.3[16] using a grid spacing of 0.5 Å and a surface probe of 0.8 Å. 'Dummy waters' were selected manually and then all others within 1.8 Å of seed waters were selected to create the surface visualised in the final figures. Analysis of channels was cross-referenced with existing biochemical data and surface charge characteristics of channels where available.
